# SH2D2A is an indicator of favourable prognosis in bladder cancer and is enriched in activated Treg cells

**DOI:** 10.1101/2025.01.13.632739

**Authors:** Brian C. Gilmour, Johan Georg Visser, Alvaro Köhn-Luque, Paweł Borowicz, Andreas Lossius, Anne Spurkland

## Abstract

Given the expanding availability of RNA-seq and other such high dimensional data it is now possible to consider a complementary approach working in the opposite direction *i*.*e*., from transcriptomics up towards protein function. This approach may prove fruitful in producing information on the function of cytosolic-bound proteins such as adapter proteins. To test the validity of this method, we made use of several public datasets to interrogate the cancer-specific role of the adapter protein SH2D2A: a protein enriched in T and NK cells and a known interactor of the kinase LCK, whose function remains uncertain. We found that SH2D2A is a favourable marker for prognosis in urothelial bladder cancer (BLCA). Digging further, we identified a population of SH2D2A^+^ FOXP3^+^ IL2RA^hi^ activated Tregs as the main expressors of SH2D2A in BLCA. This suggests that the expression of SH2D2A in these Tregs contributes to a beneficial prognostic effect. Further comprehension of SH2D2A’s function in these cells holds the potential for advancing treatment in BLCA and diversifying possible targets for immunotherapies.

---

The function of many proteins, including adapter proteins, remains largely unknown. These proteins lack direct enzymatic activity and act as molecular bridges or scaffolds in larger signalling processes^1,2^, such proteins can have a strong impact on signalling in immune cells, and thus understanding the way they variously impact signalling in immune cells can prove an important tool in the current expansion and diversification of immunotherapies.

SH2D2A, also known as T cell specific adapter protein (TSAd), is one such adapter protein, and is specifically enriched in T and NK cells, becoming further expressed upon activation^3-5^. SH2D2A is thought to be an adapter for various immune kinases^6-13^. However, despite the demonstrated ability of SH2D2A and LCK to interact, removal of the former does not appear to strongly impact TCR signalling. Likewise, SH2D2A knockout mice show only subtle changes in immune system function^12-14^. As such, SH2D2A remains for the time being one of many proteins whose function, despite research to elucidate it, remains largely unknown.

As SH2D2A is preferentially expressed in immune cells, while also being identified as an important adapter in epithelial cells, we reasoned that the presence or lack of SH2D2A may play a role in cancer survival. Prior research has already suggested that TCR-transgenic mice lacking SH2D2A have improved resistance to myeloma^15^, while SH2D2A deficient mice also grow smaller experimental sarcomas^7,16^. To explore this notion further, we turned to public datasets of bulk and single cell (sc)-RNA-seq data of cancer patients.

Here we identified that SH2D2A is a negative prognostic factor for glioblastoma (GBM), uterine corpus endometrial cancer (UCEC), kidney papillary cell cancer (KIRP), and kidney clear cell cancer (KIRC), while also observing that SH2D2A is a positive prognostic factor in bladder urothelial cancer (BLCA). We determined that the main contributor of SH2D2A expression in tumours is T cells. Exploring the effects of SH2D2A in bladder cancer further, we found that an expanded cluster of activated CD4^+^ cells, including a notable enrichment of activated FOXP3 Treg cells, was the largest contributor to SH2D2A expression in the bladder tumour microenvironment. This suggests that the CD4^+^ FOXP3^+^ SH2D2A^+^ population may be the root cause of the SH2D2A-associated positive effect on prognosis.

## Methods

### Sourced patient data

This study sources data from several publicly available datasets, as detailed below. Consent was obtained for all patients included in each of the referenced datasets.

### Processing & analysis of the TCGA cancer patient data

Bulk RNA-seq fragments per kilobase million (FPKM) values from the TCGA were recompiled into a matrix of Ensembl gene IDs and sample IDs for each constituent cancer type (**Table 1**). A pseudo-single cell object (the “single tumour” (st)-RNA-seq object) was compiled by treating each individual sample ID entry as a “cell” – the resultant object was then analysed using the R package Seurat. The R package MiloR was then used to determine the common genes over- and under-expressed by the SH2D2A-enriched sample neighbourhoods, as described elsewhere^17^, gene ontology (GO) analysis was then done using the package clusterProfiler^18^.

**TABLE 1.** The Cancer Genome Atlas’s cancer IDs, their full names, and the names used in this study.

| TCGA name | ID | Form in this paper |
| --- | --- | --- |
| Bladder urothelial carcinoma | BLCA | Bladder, urothelial |
| Breast invasive carcinoma | BRCA | Breast, invasive |
| Cervical and endocervical cancers | CESC | Cervical & endocervical |
| Colon adenocarcinoma | COAD | Colon |
| Glioblastoma multiforme | GBM | Glioblastoma multiforme |
| Head and neck squamous cell carcinoma | HNSC | Head & neck, squamous cell |
| Kidney chromophobe | KICH | Kidney, chromophobe |
| Kidney renal clear cell carcinoma | KIRC | Kidney, clear cell |
| Kidney renal papillary cell carcinoma | KIRP | Kidney, papillary cell |
| Liver hepatocellular carcinoma | LIHC | Liver |
| Lung adenocarcinoma | LUAD | Lung |
| Lung squamous cell carcinoma | LUSC | Lung, squamous cell |
| Ovarian serous cystadenocarcinoma | OV | Ovarian |
| Pancreatic adenocarcinoma | PAAD | Pancreatic |
| Prostate adenocarcinoma | PRAD | Prostate |
| Rectum adenocarcinoma | READ | Rectum |
| Skin cutaneous melanoma | SKCM | Melanoma |
| Stomach adenocarcinoma | STAD | Stomach |
| Testicular germ cell tumors | TGCT | Testicular, germ cell |
| Thyroid carcinoma | THCA | Thyroid |
| Uterine corpus endometrial carcinoma | UCEC | Uterine corpus, endometrial |

### Analysis of IMvigor210 data

The RNA expression matrix and meta data for patients from the IMvigor210 study were loaded into R as described^19^. Patient samples were classified based on the origin of the biopsy, as well as split into groups based on whether they expressed SH2D2A in amounts higher or lower than average, and metadata summarised per each group (**Table S2**).

### Survival curve analysis of TCGA and IMvigor210 data

Survival curves were created using the R package survival for both the TCGA and IMvigor210 bulk RNA datasets. For the TCGA data, the population was divided into a high or low SH2D2A-expressing cohort based on the average expression of SH2D2A in the given cancer type. Survival analysis was then carried out between these two groups and used for generation of Kaplan-Meier survival plots for each type of cancer. For the IMvigor210 data, survival plots were created for the entire cohort, and for the samples originating only from the bladder organ, with the average expression of SH2D2A based on those of the patients in each cohort.

### Cell type assignment and ADT assay import via Azimuth

For the scRNA-seq datasets GSE190888, GSE149652, as well as the Tabula Sapiens Project’s reference dataset for the bladder, immune cell types were annotated using the R package Azimuth and the peripheral blood mononuclear cell (PBMC) reference dataset *pbmcref*^20^. *Azimuth* was also used concomitantly to import and assign antibody-derived tag (ADT, CiteSeq)^21^ data from a TotalSeq™ antibody assay of 228 immune-relevant surface proteins^20^.

### Analysis of GSE190888

The single-cell transcriptome dataset GSE190888^22^ from bladder cancer was downloaded from GEO and the expression matrix annotated with downloaded Ensembl gene IDs and patient IDs. No metadata file was immediately available, and thus cursory metadata was generated from GSE190888’s sample entries GSM5733425, GSM5733426, GSM5733427, and GSM5733428 available on the GEO Accession page. The expression matrix and metadata were combined as a Seurat object and processed as per conventional analysis using Seurat as described above and dimensional reduction displayed using uniform manifold approximation and projection (UMAP). SH2D2A-expressing clusters where then extracted and the new Seurat object processed again as above, with the dimensional reduction this time visualised using *t*-distributed stochastic neighbour embedding (*t*-SNE). Immune cell types were assigned, and the ADT assay imported using *Azimuth* as described above. The proportions of immune cell types and their SH2D2A expression were summarised for each condition, *i*.*e*. primary BLCA, recurrent BLCA, and cystitis glandularis (C. glandularis). Differentially regulated elements (RNA and ADT) in the SH2D2A-enriched cell types were then found and GO analysis performed, with the top upregulated genes and their corresponding upregulated GO terms plotted as a chord plot using the R package GOplot^23^.

### Combined analysis of T cells from GSE149652 and the Tabula Sapiens’s bladder reference dataset

The dataset GSE149652^24^, consisting of T cells from BLCA patients undergoing αPD-L1 therapy, chemotherapy, or from untreated patients, was downloaded from GEO and the expression matrix annotated with downloaded Ensembl gene IDs and patient IDs. A Seurat object was then created from the annotated expression matrix and the downloaded patient metadata. To provide a reference for T cells from the non-malignant bladder, GSE149652 was combined with the T cells from the Tabula Sapiens Project’s reference dataset for the bladder using *Seurat*’s merge function. The combined Seurat object was then processed as per conventional analysis using *Seurat* as described above and dimensional reduction visualised using t-SNE to enhance cluster borders. Immune cell types were then assigned, and the ADT assay imported using Azimuth as described above. Any cells not annotated as T cells were removed to avoid ambiguity. Clusters enriched for Tregs and SH2D2A were identified, and cluster markers were determined for the main SH2D2A-expressing Treg-enriched cluster, cluster 7. All Treg- and SH2D2A-enriched clusters were then used for side-by-side GO analysis, highlighting commonalities between clusters 7 and 8, which were then linked to the manual annotation IL2RA^hi^ present from the original analysis by Oh *et al*.^24^, markers of which were then used for further survival analysis using bladder organ-derived cancers samples from IMvigor210^19^.

## Code availability

All analytical work was carried out using pre-existing algorithms and packages, and no novel algorithms or packages were developed for the purpose of this study. Scripts used for bioinformatic analysis will be made available on GitHub with the published article.

## Data availability

No new datasets were generated by this study. A summary of the datasets used in this study and their sources is given in Table S1.

## Results

### Higher SH2D2A expression is a prognostic factor in five types of cancer

To assess the correlation of SH2D2A with cancer prognosis, we analyzed the TCGA dataset of RNA-seq data from 21 cancer types (**Table 1,** sample numbers in **Figure 1A**). We split the data by cancer type and isolated the FPKM values of SH2D2A (summarized in **Figure 1B**). We then plotted SH2D2A expression against normalized survival time, observing a general tendency for survival time to decrease with increased SH2D2A expression (**Figure 1C**). Each cancer type was individually analyzed by splitting patients into high (SH2D2A > average) and low (SH2D2A ≤ average) SH2D2A subgroups. Most cancer types showed no significant difference in survival (**Figure S1**), but high SH2D2A favored survival in BLCA (p = 0.015, **Figure 1D**) and was linked to lower survival in GBM (p = 0.0068), UCEC (p = 0.01), KIRP (p = 0.029), and KIRC (p = 0.00022, **Figure 1E**).

**Figure 1.**
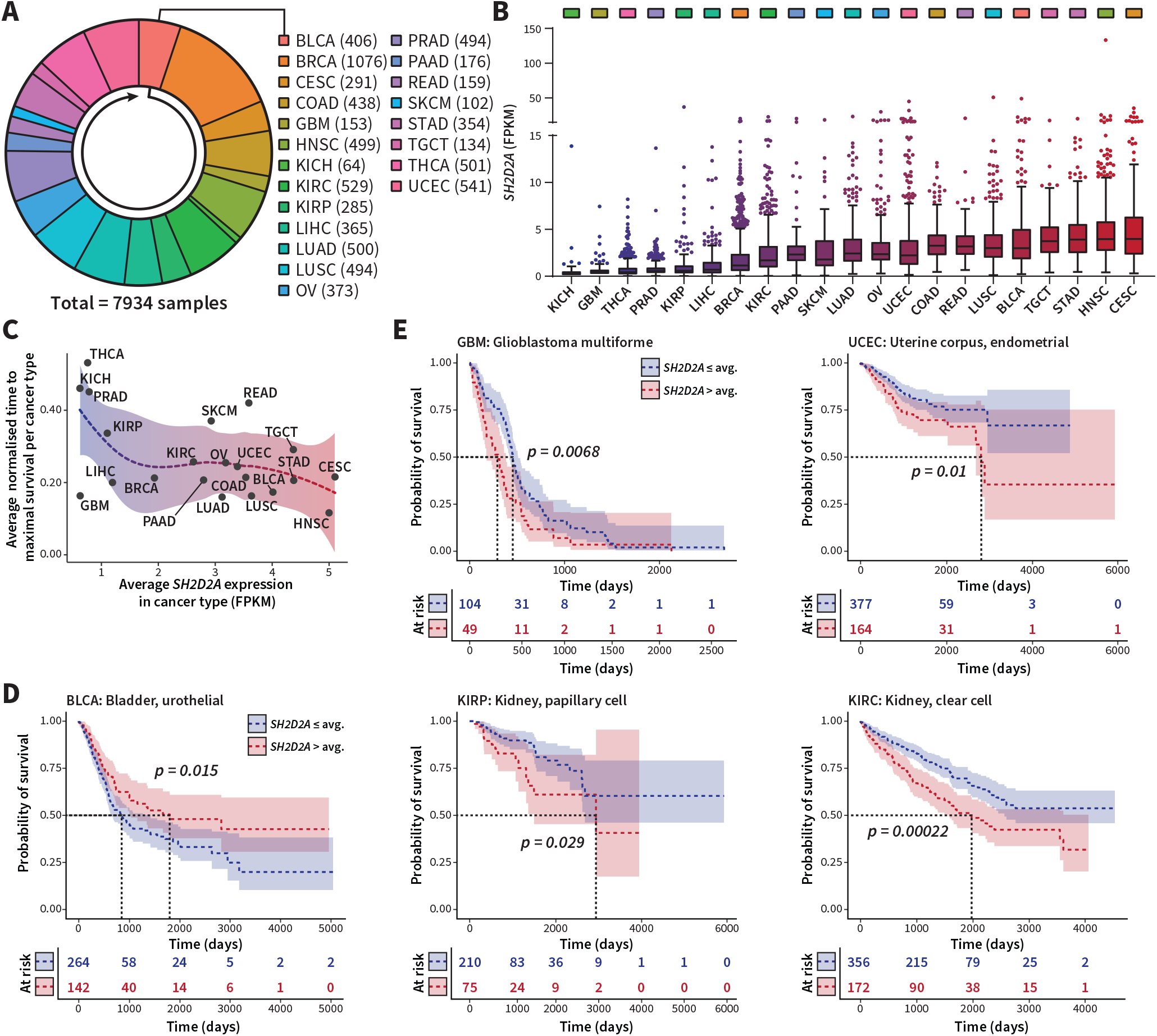
SH2D2A is a prognostic factor in BLCA, GBM, UCEC, KIRP and KIRC. **(A)** Number of patient samples in each cancer subtype in the TCGA dataset, ordered clockwise alphabetically. **(B)** Fragments per kilobase million (FPKM) SH2D2A expressed in each cancer subtype plotted as box-and-whisker plots. Tumour subtypes are ordered left-to-right in order of decreasing median FPKM of SH2D2A. **(C)** Relation between average normalised time to maximal survival for each cancer type to the average SH2D2A expression in that cancer type, plotted with logarithmic regression line and 95% confidence interval. **(D-E)** Kaplan-Meier survival curves of SH2D2A > avg. and SH2D2A ≤ avg. groups showing a beneficial effect of SH2D2A on prognosis in BLCA **(D)** and a detrimental effect of SH2D2A on prognosis in GBM, UCEC, KIRP and KIRC **(E)**.

### SH2D2A expression in tumours is associated with higher expression of immune genes

We then investigated whether the prognostic effect of SH2D2A arose from immune cells, endothelia, or the tumors themselves by compiling TCGA’s bulk RNA-seq data into a single tumor RNA-seq object using Seurat (**Figure 2A**). Dimensional reduction using UMAP showed SH2D2A abundance in various squamous cell cancers (**Figure 2B**). Using *MiloR*, we divided tumors into neighborhoods characterized by SH2D2A levels (**Figure 2B**, lower inset). Differential gene expression analysis between SH2D2A-abundant and SH2D2A- sparse neighborhoods (**Figure 2C**) showed enrichment of immune-related processes (**Figure 2D**), suggesting a link between high SH2D2A and activated immune cells in the tumor environment.

**Figure 2.**
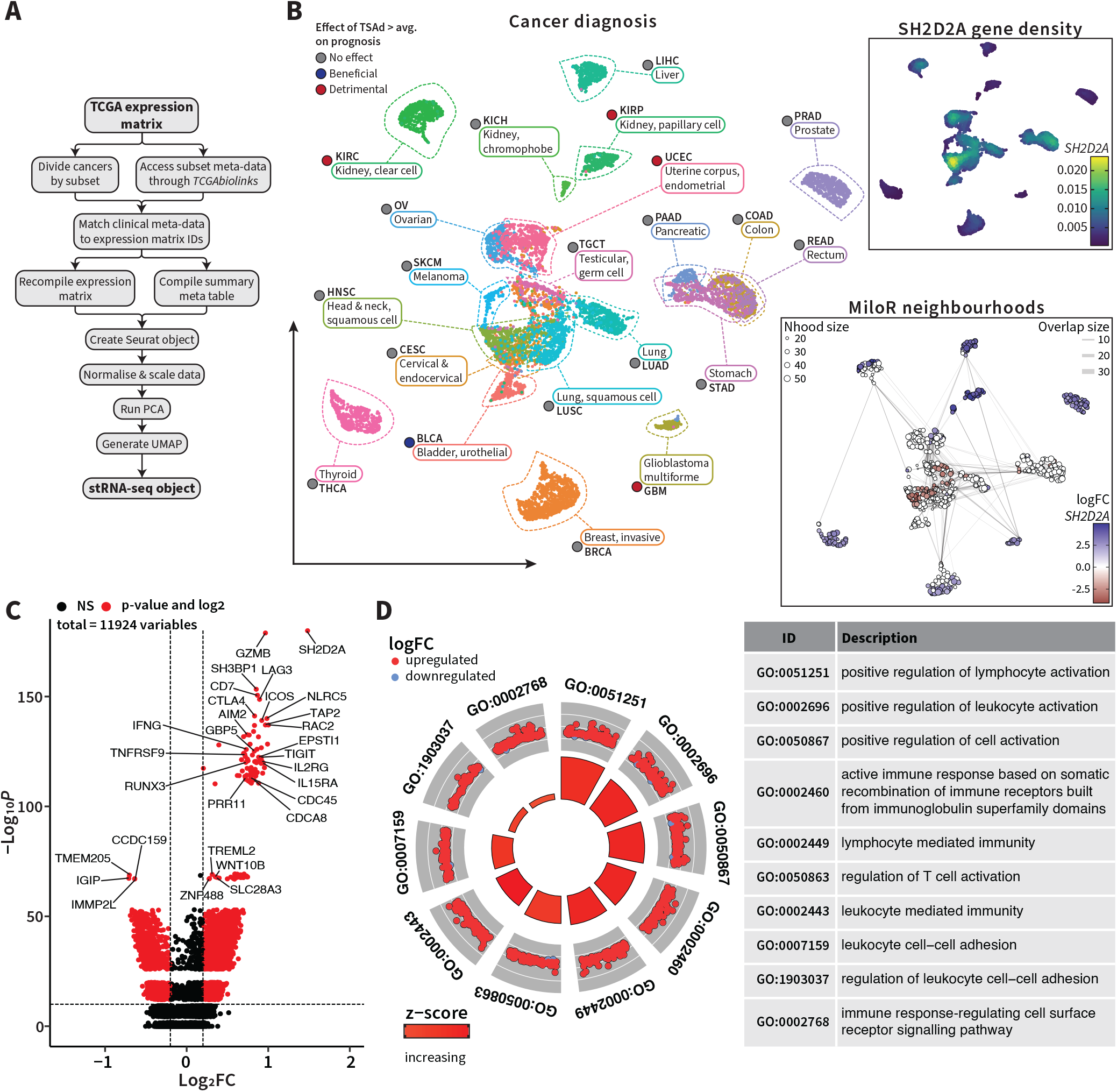
Higher SH2D2A expression in tumours is linked to expression of immune genes. **(A)** Workflow summary to generate the “single tumour” RNA-seq object for analysis in **Seurat. (B)** UMAP of the ***“single tumour”*** RNA-seq object, coloured and annotated to show the locations of the different cancer subtypes, as well as the effect of SH2D2A > avg. on prognosis in each subtype, SH2D2A density was mapped onto the UMAP **(B, *upper inset*)** as well as the neighbourhoods as generated by **MiloR (B, *lower inset*). (C)** Volcano plot of differentially expressed genes favourable expressed with SH2D2A (right) and favourably expressed in SH2D2A’s absence (left). **(D)** Circle plot of GO pathway analysis, showing GO terms enriched for associated upregulated and downregulated genes.

### SH2D2A expression in BLCA does not predict response to ICI therapy

Given the association between SH2D2A and active immune responses, we examined SH2D2A as a marker for anti-cancer immune responses in BLCA using the IMvigor210 trial data of atezolizumab (αPD-L1) therapy. Patients were divided into SH2D2A above- and below-average subgroups. While the higher SH2D2A subgroup showed only a trend towards better survival in the full dataset (p = 0.13, **Figure 3A**), significant beneficial effects were noted when focusing on only bladder-derived samples (p = 0.01, **Figure 3B**). SH2D2A > avg. tumors were more often categorized as “Genomically unstable” or “SCC-like,” with more inflamed characteristics and higher immune cell infiltration (**Figure 3C**). Transcriptomic analysis of the 75 most consistently upregulated genes in the SH2D2A > avg. group confirmed a higher degree of immune cell presence in SH2D2A > avg. tumors (**Figure 3D**), supported by GO pathway analysis (Figure S2).

**Figure 3.**
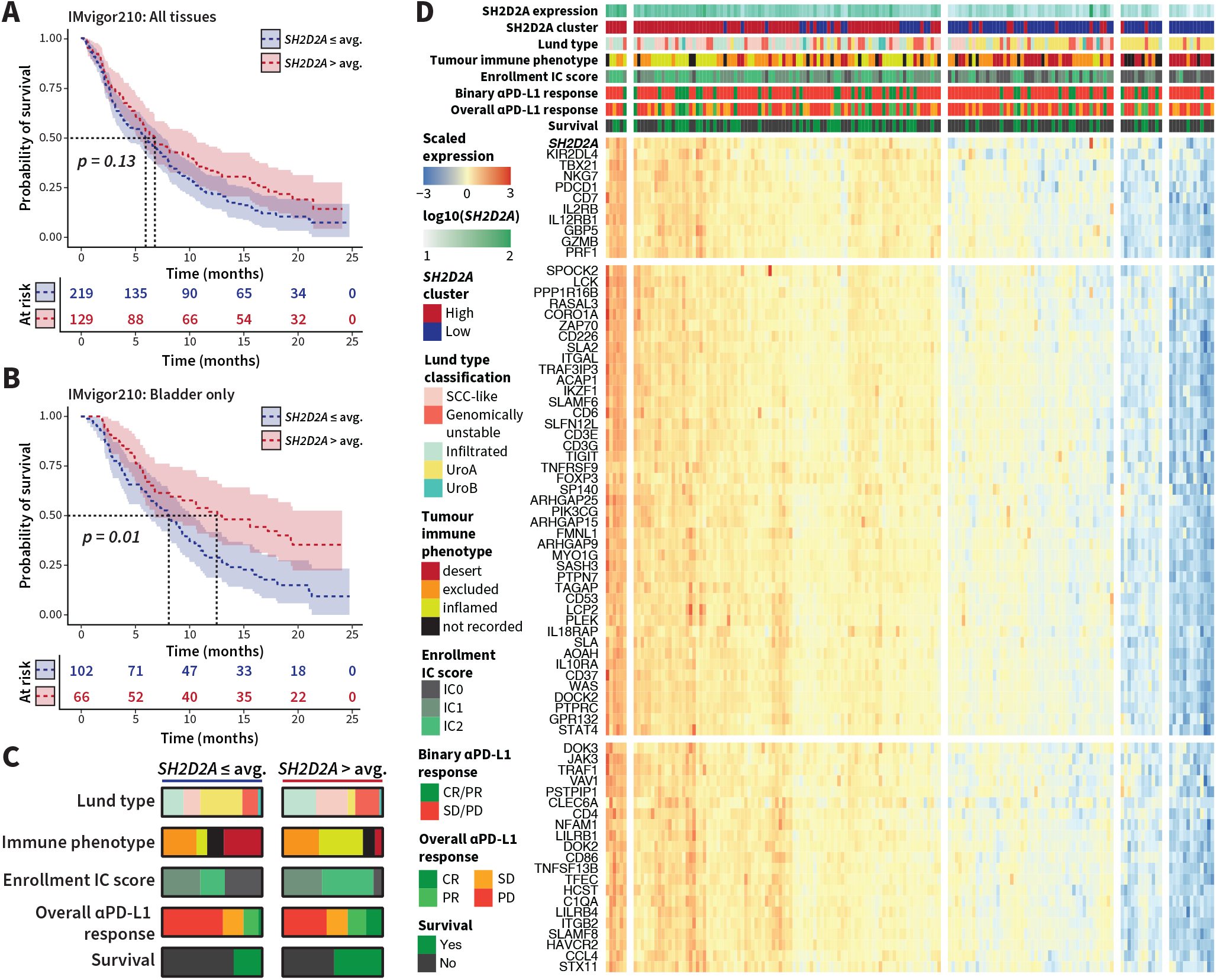
SH2D2A is not a predictor of response to αPD-L1 therapy in BLCA. **(A-B)** Kaplan-Meier survival curves for IMvigor210 patients in SH2D2A > avg. and SH2D2A ≤ avg. groupings with tumour samples deriving from all sampled tissues in IMvigor210 (bladder + metastases) **(A)** and from the bladder organ only **(B). (C)** Summary comparison of Lund type, immune phenotype, IC score at enrollment, overall αPD-L1 therapy response, and survival metadata descriptors for the SH2D2A > avg. and SH2D2A ≤ avg. groupings (see Table S2 for descriptions of IMvigor210 metadata contents used in this paper). **(D)** Heatmap of the top 75 upregulated genes in the SH2D2A avg. group, as determined by Z-normalisation of each gene across all patients in the group.

### Tregs are enriched for SH2D2A in the BLCA bladder

To identify immune cells responsible for SH2D2A expression in BLCA, we analyzed scRNA-seq data (GSE190888) and confirmed that SH2D2A was expressed primarily in immune cells (Figure S3). We gated out the SH2D2A-expressing cells (**Figure 3A**) and, using Azimuth, we annotated them using the PBMC reference dataset *pbmcref*^20^, concomitantly importing an ADT assay of 228 immune-relevant cell surface proteins onto the cells, and cell identities visualised via *t*-SNE plot (**Figure 3B**). This revealed Tregs as highly enriched for SH2D2A (**Figure 4D,** red box). Density mapping of SH2D2A and FOXP3 showed substantial overlap (**Figure 4E**), and transcriptional markers along with enriched GO pathways of the SH2D2A-expressing cells suggested they were dominated by active, tolerance-promoting Tregs (**Figure 4F**).

**Figure 4.**
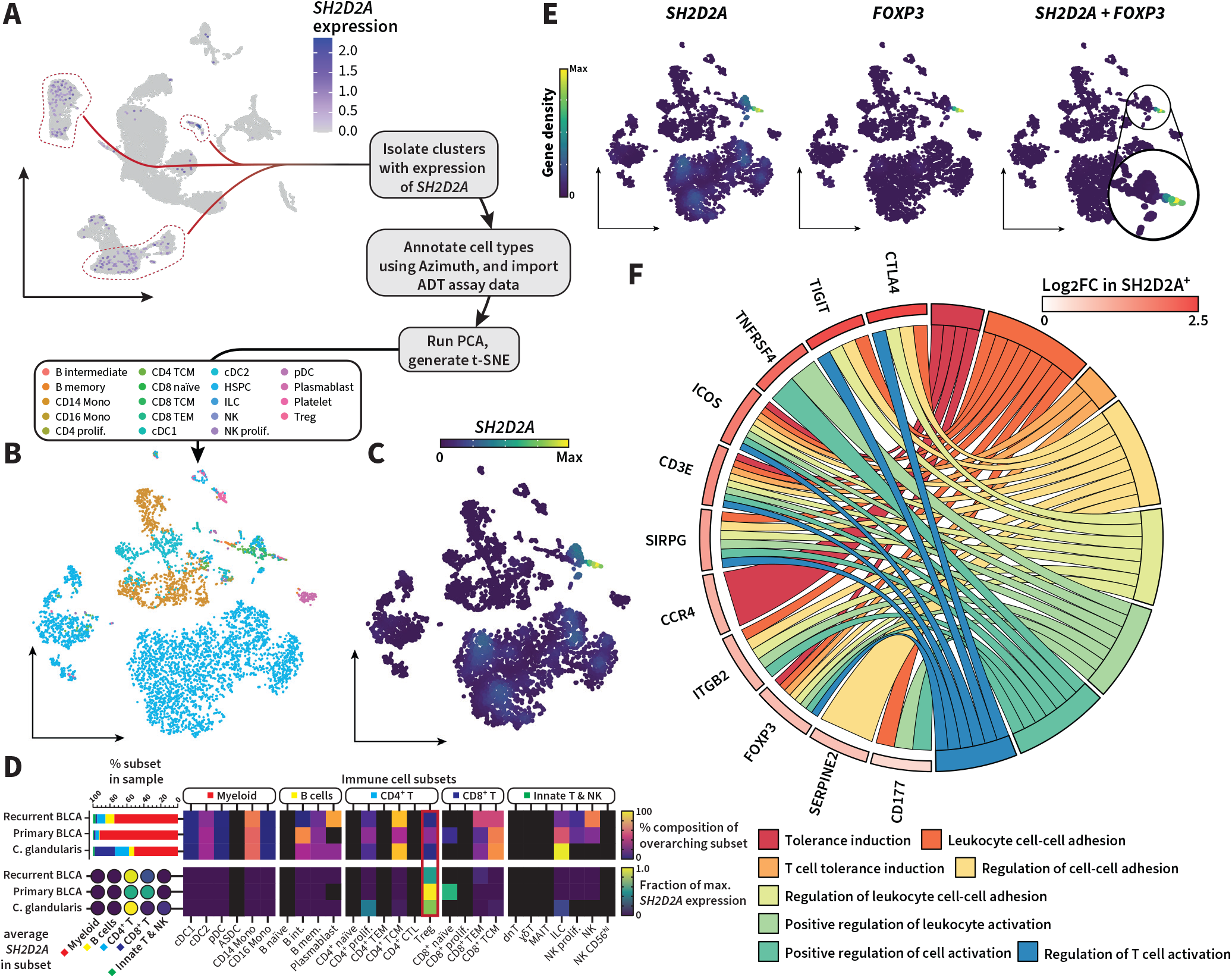
Tregs are enriched for SH2D2A in the BLCA bladder. **(A)** UMAP of GSE190888 mapped with SH2D2A expression levels, red lines indicate clusters isolated for further analysis, with the processing workflow for the isolated cells also depicted. **(B)** *t*-SNE of SH2D2A^+^ cells coloured by cell type mapping output from **Azimuth. (C)** Density plot of SH2D2A on the *t*-SNE plot from **(B). (D)** Summary descriptions per each condition present in GSE190888 *i*.*e*., primary BLCA, recurrent BLCA, and cystitis glandularis (C. glandularis) of cell type abundance (***upper***) and SH2D2A abundance (***lower***) in the overarching cell subtypes (***left***) and in the cell types themselves (***right***), the entry for Tregs, which are enriched for SH2D2A, is highlighted (***red box***). **(E)** Density plots of SH2D2A, FOXP3 and a joint density plot of SH2D2A and FOXP3 mapped onto the *t-*SNE of **(B). (F)** Chord plot of the gene markers of the entire population of SH2D2A^+^ cells in **(B-C)**, their log_2_ fold change, and the GO terms enriched in the SH2D2A^+^ cell population that each gene is connected to.

### SH2D2A is enriched in an activated population of Treg cells in BLCA

Given the low Treg count in GSE190888, we sought another BLCA dataset (GSE149652) with more immune cells, including CD4^+^ and CD8^+^ T cells from BLCA patients undergoing various forms of therapy, or taken from patients prior to treatment. We merged this dataset with T cells from the Tabula Sapiens healthy reference dataset for the bladder. Dimensional reduction of the merged datasets identified 27 clusters (**Figure 5A**), with some clusters dominated by a single patient (**Figure 5B**) and an overall expression of SH2D2A (**Figure 5C**). Many high-expressing clusters consisted primarily of innate T or T effector cells (**Figure 5D**), with clusters 7 and 8 showing a higher abundance of Treg cells (**Figure 5E**). Overlaps between SH2D2A and Treg marker FOXP3 were particularly strong in both these clusters (**Figure 5F**). Differential expression and GO enrichment analyses once again indicated an activated Treg state (**Figure 5G-H**), as contrasted with other clusters containing Tregs. Both clusters 7 and 8 also contributed an exaggerated proportion of the SH2D2A in a Treg subset annotated as IL2RA^hi^ by Guo *et al*. (**Figure 5I,** full list of manually annotated cell types in Table S3) characterised as expressing several-fold higher amounts of IL2RA, FOXP3, TIGIT, TNFRSF4/9/18 and CD27 transcripts^24^.

**Figure 5.**
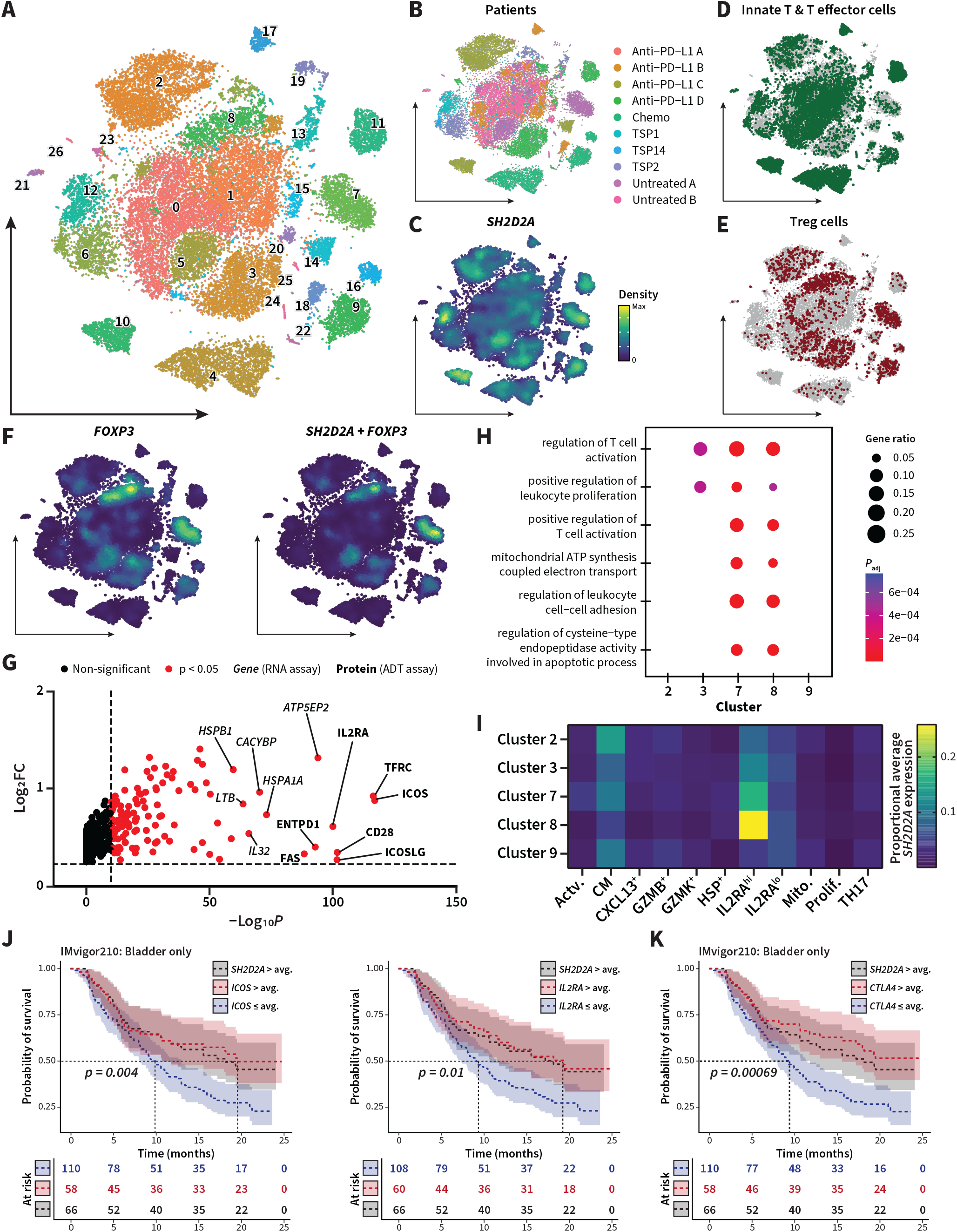
SH2D2A is enriched in an activated subset of Treg cells in BLCA. **(A)** *t-*SNE plot of combined dataset consisting of GSE149652 and T cells from the Tabula Sapiens bladder reference dataset, clusters derived from unsupervised clustering are shown. **(B-E)** *t-*SNE of the combined dataset mapped for patient of origin **(B)**, density of *SH2D2A* expression **(C)**, cells annotated by Azimuth as innate T or effector T cells **(D)**, and cells annotated by Azimuth as Treg cells **(E). (F)** Density plot of *FOXP3* expression and joint density plot of *SH2D2A* and *FOXP3*. **(G)** Volcano plot of upregulated genes (*italic*, RNA assay) and extrapolated surface protein values (**bold**, ADT assay) of cluster 7, showing only upregulated elements. **(H)** Clustered GO analysis between clusters 2, 3 and 7-9. **(I)** Average SH2D2A expression in each cell type annotation derived from Oh *et al*.^24^ in clusters 2, 3 and 7-9, (…) (…) average SH2D2A expression is scaled to the proportion of each cell type annotation in each cluster. **(J-K)** Kaplan-Meier survival curves for IMvigor210 patients with tumours from the bladder organ in above/below-or-equal groupings for ICOS/IL2RA **(J)** and CTLA4 **(K)**, with the survival curve for the above-average grouping for SH2D2A shown for comparison (**black**).

To further test the overlap of clusters 7 and 8 with this IL2RA^hi^ Treg subset, we ran survival analyses with the IMvigor210 dataset with ICOS and IL2RA (**Figure 5J**), two previously identified markers of this Treg population, with both providing a similar beneficial effect on survival as SH2D2A (p_ICOS_ = 0.004, p_IL2RA_ = 0.01). CTLA4, another Treg marker, also correlated with higher survival, comparable to SH2D2A (**Figure 5K**), further suggesting that it is this population of SH2D2A^+^ Tregs that is providing a beneficial effect on prognosis in BLCA.

## Discussion

Analysing publicly available datasets, we here identified SH2D2A as a beneficial prognostic factor in BLCA. ICOS^+^ FOXP3^+^ IL2RA^hi^ Treg cells were the main contributors to SH2D2A expression in BLCA tumours. These SH2D2A- expressing ICOS^+^ FOXP3^+^ IL2RA^hi^ cells feature a noted enrichment of Tregs annotated by Oh *et al*. as CD4_IL2RAHI_^24^ (glossed IL2RA^hi^ here). Furthermore, TCR sequencing by Oh *et al*. revealed these cells as being significantly expanded and expressing private TCR repertoires not shared with other CD4^+^ T cell populations, thus unlikely to be induced Tregs^24^.

Similar populations of non-induced Tregs expressing high amounts of FOXP3 and IL2RA have likewise been reported in non-small cell lung cancer^29^ and melanoma^30^ as well as cancers of the breast^31^, colon^31^, lung^32^, and liver^33^. However, in these tissues, this population is associated with worse outcome. This is backed up by our own analysis of the IMvigor210 cohort^19^, where a positive correlation of SH2D2A to improved prognosis appears to be limited to the bladder organ itself and does not apply to BLCA-derived metastases in other tissues.

The specific effects of SH2D2A expression and its consequences in Treg cells is understudied. SH2D2A knockout mice have been shown to be more prone to graft rejection, a phenomenon that was linked to a reduced ability on the part of Tregs to promote graft tolerance^34^. In contrast, effector responses in the SH2D2A knockout mice were not affected.

The current understanding of Tregs in the immune response against cancer is that they inhibit the anti-tumor response of other immune cells^35,36^. It is thus tempting to assume that the beneficial effect of SH2D2A on prognosis in BLCA arises from the modulation of Tregs, such that their ability to inhibit ongoing immune responses is diminished. The notion that SH2D2A modulates T cell responses in cancer is backed up by preliminary analyses similar to that done for BLCA in this study, where we have observed that in the cancer subtypes where high SH2D2A was correlated with a worse prognosis, *i*.*e*. GBM, UCEC, KIRP and KIRC, expression of SH2D2A was far more disperse, and includes a higher degree of expression in CD8^+^ T cells, innate-like T cells, and NK cells (Figure S4). Likewise, the main CD8^+^ expressor in BLCA as annotated by Oh *et al*. (CD8_ENTPD1_) is a group of tumour-reactive T effector cells expressing several markers of exhaustion, such as HAVCR2^37-40^, LAG3^41,42^, and CXCL13^30,33,43^.

Thus, SH2D2A may be a marker of inhibited immune cell function or may play a more direct role in inhibitory signaling pathways in immune cells. However, SH2D2A is expressed to some degree in all of the 21 cancer subtypes available for analysis from TCGA, yet a differential prognostic effect associated with a higher presence of SH2D2A was only visible in BLCA, GBM, UCEC, KIRP, and KIRC.

The expression profile both at the transcript level and the inferred surface protein level, suggests that the SH2D2A^+^ Treg cells in BLCA are primed and actively inducing tolerance, thus impeding ongoing immune reactions. It is possible that the bladder mucosa represents a unique immune environment45, where SH2D2A^+^ Tregs may play a paradoxical effect on BLCA progression or response to therapy.

As such, more research is needed to uncover the actual function of SH2D2A in Tregs in BLCA, and what cellular state its expression is indicating. The identification of this function, the receptor or signalling pathway it falls under, or the overall cellular state as indicated by SH2D2A, will offer new targets for the future treatment of BLCA, that can perhaps be expanded further to GBM, UCEC, KIRP, and KIRC, and add to the arsenal of targets for immunotherapies.

## Abbreviations

BLCA: bladder urothelial cancer
C. glandularis: cystitis glandularis
DEA: differential expression analysis
FPKM: fragments per kilobase million
GBM: glioblastoma multiforme
GEO: Gene Expression Omnibus
GO: gene ontology
IC: immune cell
ICI: immune checkpoint inhibitor
KIRC: kidney clear cell cancer
KIRP: kidney papillary cell cancer
NK: natural killer
PBMC: peripheral blood mononuclear cell
PCA: principal component analysis
(sc)RNA-seq: (single cell) RNA
sequencing
SCID: severe combined immunodeficiency
SEM: standard error of the mean
TCGA: the Cancer Genome Atlas
TCR: T cell receptor
Treg: T regulatory cell
t-SNE: t-distributed stochastic neighbour embedding
UCEC: uterine corpus
endometrial cancer
UMAP: uniform manifold approximation and projection
VEGF: vascular endothelial growth factor

## Author contributions

Brian C. Gilmour: Conceptualisation, Methodology, Software, Investigation, Formal Analysis, Data Curation, Visualisation, Writing – Original Draft, Writing – Review & Editing. Johan Georg Visser: Conceptualisation, Methodology, Writing – Review & Editing. Alvaro Köhn-Luque: Validation, Supervision. Andreas Lossius: Resources. Paweł Borowicz: Conceptualisation, Methodology. Anne Spurkland: Conceptualisation, Supervision, Project Administration, Funding Acquisition, Writing – Review & Editing

## Acknowledgements

Funding: This work was supported by a grant from the Norwegian Research Council (#302647).

## References

1. Borowicz, P., Chan, H., Hauge, A., and Spurkland, A. (2020). Adaptor proteins: Flexible and dynamic modulators of immune cell signalling. Scandinavian Journal of Immunology 92, e12951. 10.1111/sji.12951.

2. Flynn, D.C. (2001). Adaptor proteins. Oncogene 20, 6270–6272. 10.1038/sj.onc.1204769.

3. Kolltveit, K.M., Granum, S., Aasheim, H.C., Forsbring, M., Sundvold-Gjerstad, V., Dai, K.Z., Molberg, O., Schjetne, K.W., Bogen, B., Shapiro, V.S., et al. (2008). Expression of SH2D2A in T-cells is regulated both at the transcriptional and translational level. Mol Immunol 45, 2380–2390. 10.1016/j.molimm.2007.11.005.

4. Dai, K.Z., Johansen, F.E., Kolltveit, K.M., Aasheim, H.C., Dembic, Z., Vartdal, F., and Spurkland, A. (2004). Transcriptional activation of the SH2D2A gene is dependent on a cyclic adenosine 5’-monophosphate-responsive element in the proximal SH2D2A promoter. J Immunol 172, 6144–6151. 10.4049/jimmunol.172.10.6144.

5. Sundvold, V., Torgersen, K.M., Post, N.H., Marti, F., King, P.D., Røttingen, J.A., Spurkland, A., and Lea, T. (2000). T cell-specific adapter protein inhibits T cell activation by modulating Lck activity. J Immunol 165, 2927–2931. 10.4049/jimmunol.165.6.2927.

6. Sun, Z., Li, X., Massena, S., Kutschera, S., Padhan, N., Gualandi, L., Sundvold-Gjerstad, V., Gustafsson, K., Choy, W.W., Zang, G., et al. (2012). VEGFR2 induces c-Src signaling and vascular permeability in vivo via the adaptor protein TSAd. J Exp Med 209, 1363–1377. 10.1084/jem.20111343.

7. Matsumoto, T., Bohman, S., Dixelius, J., Berge, T., Dimberg, A., Magnusson, P., Wang, L., Wikner, C., Qi, J.H., Wernstedt, C., et al. (2005). VEGF receptor-2 Y951 signaling and a role for the adapter molecule TSAd in tumor angiogenesis. Embo j 24, 2342–2353. 10.1038/sj.em-boj.7600709.

8. Granum, S., Andersen, T.C., Sørlie, M., Jørgensen, M., Koll, L., Berge, T., Lea, T., Fleckenstein, B., Spurkland, A., and Sundvold-Gjerstad, V. (2008). Modulation of Lck function through multisite docking to T cell-specific adapter protein. J Biol Chem 283, 21909–21919. 10.1074/jbc.M800871200.

9. Granum, S., Sundvold-Gjerstad, V., Dai, K.Z., Kolltveit, K.M., Hildebrand, K., Huitfeldt, H.S., Lea, T., and Spurkland, A. (2006). Structure function analysis of SH2D2A isoforms expressed in T cells reveals a crucial role for the proline rich region encoded by SH2D2A exon 7. BMC Immunol 7, 15. 10.1186/1471-2172-7-15.

10. Sundvold-Gjerstad, V., Granum, S., Mustelin, T., Andersen, T.C., Berge, T., Shapiro, M.J., Shapiro, V.S., Spurkland, A., and Lea, T. (2005). The C terminus of T cell-specific adapter protein (TSAd) is necessary for TSAd-mediated inhibition of Lck activity. Eur J Immunol 35, 1612–1620. 10.1002/eji.200425638.

11. Choi, Y.B., Kim, C.K., and Yun, Y. (1999). Lad, an adapter protein interacting with the SH2 domain of p56lck, is required for T cell activation. J Immunol 163, 5242–5249.

12. Rajagopal, K., Sommers, C.L., Decker, D.C., Mitchell, E.O., Korthauer, U., Sperling, A.I., Kozak, C.A., Love, P.E., and Bluestone, J.A. (1999). RIBP, a novel Rlk/ Txk- and itk-binding adaptor protein that regulates T cell activation. J Exp Med 190, 1657–1668. 10.1084/jem.190.11.1657.

13. Berge, T., Sundvold-Gjerstad, V., Granum, S., Andersen, T.C.B., Holthe, G.B., Claesson-Welsh, L., Andreotti, A.H., Inngjerdingen, M., and Spurkland, A. (2010). T Cell Specific Adapter Protein (TSAd) Interacts with Tec Kinase ITK to Promote CXCL12 Induced Migration of Human and Murine T Cells. PLOS ONE 5, e9761. 10.1371/journal.pone.0009761.

14. Marti, F., Garcia, G.G., Lapinski, P.E., MacGregor, J.N., and King, P.D. (2006). Essential role of the T cell–specific adapter protein in the activation of LCK in peripheral T cells. Journal of Experimental Medicine 203, 281–287. 10.1084/jem.20051637.

15. Berge, T., Grønningsæter, I.H.B., Lorvik, K.B., Abrahamsen, G., Granum, S., Sundvold-Gjerstad, V., Corthay, A., Bogen, B., and Spurkland, A. (2012). SH2D2A Modulates T Cell Mediated Protection to a B Cell Derived Tumor in Transgenic Mice. PLOS ONE 7, e48239. 10.1371/journal.pone.0048239.

16. Wu, L.-W., Mayo, L.D., Dunbar, J.D., Kessler, K.M., Ozes, O.N., Warren, R.S., and Donner, D.B. (2000). VRAP Is an Adaptor Protein That Binds KDR, a Receptor for Vascular Endothelial Cell Growth Factor*. Journal of Biological Chemistry 275, 6059–6062. 10.1074/jbc.275.9.6059.

17. Dann, E., Henderson, N.C., Teichmann, S.A., Morgan, M.D., and Marioni, J.C. (2022). Differential abundance testing on single-cell data using k-nearest neighbor graphs. Nature Biotechnology 40, 245–253. 10.1038/s41587-021-01033-z.

18. Wu, T., Hu, E., Xu, S., Chen, M., Guo, P., Dai, Z., Feng, T., Zhou, L., Tang, W., Zhan, L., et al. (2021). clusterProfiler 4.0: A universal enrichment tool for interpreting omics data. The Innovation 2, 100141. 10.1016/j.xinn.2021.100141.

19. Mariathasan, S., Turley, S.J., Nickles, D., Castiglioni, A., Yuen, K., Wang, Y., Kadel Iii, E.E., Koeppen, H., Astarita, J.L., Cubas, R., et al. (2018). TGFβ attenuates tumour response to PD-L1 blockade by contributing to exclusion of T cells. Nature 554, 544–548. 10.1038/nature25501.

20. Hao, Y., Hao, S., Andersen-Nissen, E., Mauck, W.M., Zheng, S., Butler, A., Lee, M.J., Wilk, A.J., Darby, C., Zager, M., et al. (2021). Integrated analysis of multimodal single-cell data. Cell 184, 3573-3587.e3529. 10.1016/j.cell.2021.04.048.

21. Stoeckius, M., Hafemeister, C., Stephenson, W., Houck-Loomis, B., Chattopadhyay, P.K., Swerdlow, H., Satija, R., and Smibert, P. (2017). Simultaneous epitope and transcriptome measurement in single cells. Nature Methods 14, 865–868. 10.1038/nmeth.4380.

22. Luo, Y., Tao, T., Tao, R., Huang, G., and Wu, S. (2022). Single-Cell Transcriptome Comparison of Bladder Cancer Reveals Its Ecosystem. Frontiers in Oncology 12.

23. Walter, W., Sánchez-Cabo, F., and Ricote, M. (2015). GOplot: an R package for visually combining expression data with functional analysis. Bioinformatics 31, 2912–2914. 10.1093/bioinformatics/btv300.

24. Oh, D.Y., Kwek, S.S., Raju, S.S., Li, T., McCarthy, E., Chow, E., Aran, D., Ilano, A., Pai, C.-C.S., Rancan, C., et al. (2020). Intratumoral CD4^+^ T Cells Mediate Anti-tumor Cytotoxicity in Human Bladder Cancer. Cell 181, 1612-1625.e1613. 10.1016/j.cell.2020.05.017.

25. Lin, F., Ke, Z.-B., Xue, Y.-T., Chen, J.-Y., Cai, H., Lin, Y.-Z., Li, X.-D., Wei, Y., Xue, X.-Y., and Xu, N. (2023). A novel CD8^+^ T cell-related gene signature for predicting the prognosis and immunotherapy efficacy in bladder cancer. Inflammation Research 72, 1665–1687. 10.1007/s00011-023-01772-6.

26. Shen, C., Han, C., Li, Z., Yan, Y., Li, C., Chen, H., Fan, Z., and Hu, H. (2023). Construction and Validation of a Prognostic Model Based on Pyroptosis-related Genes in Bladder Cancer. Comb Chem High Throughput Screen. 10.2174/0113862073256363230929200157.

27. Xue, Y., Zhao, G., Pu, X., and Jiao, F. (2023). Construction of T cell exhaustion model for predicting survival and immunotherapy effect of bladder cancer based on WGCNA. Frontiers in Oncology 13.

28. Pagliarulo, F., Cheng, P.F., Brugger, L., van Dijk, N., van den Heijden, M., Levesque, M.P., Silina, K., and van den Broek, M. (2022). Molecular, Immunological, and Clinical Features Associated With Lymphoid Neogenesis in Muscle Invasive Bladder Cancer. Frontiers in Immunology 12.

29. Guo, X., Zhang, Y., Zheng, L., Zheng, C., Song, J., Zhang, Q., Kang, B., Liu, Z., Jin, L., Xing, R., et al. (2018). Global characterization of T cells in non-small-cell lung cancer by single-cell sequencing. Nature Medicine 24, 978–985. 10.1038/s41591-018-0045-3.

30. Tirosh, I., Izar, B., Prakadan, S.M., Wadsworth, M.H., Treacy, D., Trombetta, J.J., Rotem, A., Rodman, C., Lian, C., Murphy, G., et al. (2016). Dissecting the multicellular ecosystem of metastatic melanoma by single-cell RNA-seq. Science 352, 189–196. 10.1126/science.aad0501.

31. Plitas, G., Konopacki, C., Wu, K., Bos, P.D., Morrow, M., Putintseva, Ekaterina V., Chudakov, Dmitriy M., and Rudensky, Alexander Y. (2016). Regulatory T Cells Exhibit Distinct Features in Human Breast Cancer. Immunity 45, 1122–1134. 10.1016/j.immuni.2016.10.032.

32. De Simone, M., Arrigoni, A., Rossetti, G., Gruarin, P., Ranzani, V., Politano, C., Bonnal Raoul J.P., Provasi, E., Sarnicola Maria L., Panzeri, I., et al. (2016). Transcriptional Landscape of Human Tissue Lymphocytes Unveils Uniqueness of Tumor-Infiltrating T Regulatory Cells. Immunity 45, 1135–1147. 10.1016/j.immuni.2016.10.021.

33. Zheng, C., Zheng, L., Yoo, J.-K., Guo, H., Zhang, Y., Guo, X., Kang, B., Hu, R., Huang, J.Y., Zhang, Q., et al. (2017). Landscape of Infiltrating T Cells in Liver Cancer Revealed by Single-Cell Sequencing. Cell 169, 1342-1356. e1316. 10.1016/j.cell.2017.05.035.

34. Wedel, J., Stack, M.P., Seto, T., Sheehan, M.M., Flynn, E.A., Stillman, I.E., Kong, S.W., Liu, K., and Briscoe, D.M. (2019). T Cell–Specific Adaptor Protein Regulates Mitochondrial Function and CD4^+^ T Regulatory Cell Activity In Vivo following Transplantation. The Journal of Immunology 203, 2328–2338. 10.4049/jimmunol.1801604.

35. Curiel, T.J., Coukos, G., Zou, L., Alvarez, X., Cheng, P., Mottram, P., Evdemon-Hogan, M., Conejo-Garcia, J.R., Zhang, L., Burow, M., et al. (2004). Specific recruitment of regulatory T cells in ovarian carcinoma fosters immune privilege and predicts reduced survival. Nature Medicine 10, 942–949. 10.1038/nm1093.

36. Yao, W., He, J.-c., Yang, Y., Wang, J.-m., Qian, Y.-w., Yang, T., and Ji, L. (2017). The Prognostic Value of Tumor-infiltrating Lymphocytes in Hepatocellular Carcinoma: a Systematic Review and Meta-analysis. Scientific Reports 7, 7525. 10.1038/s41598-017-08128-1.

37. Beck, R.J., Sloot, S., Matsushita, H., Kakimi, K., and Beltman, J.B. (2023). Mathematical modeling identifies LAG3 and HAVCR2 as biomarkers of T cell exhaustion in melanoma. iScience 26, 106666. 10.1016/j.isci.2023.106666.

38. Fourcade, J., Sun, Z., Benallaoua, M., Guillaume, P., Luescher, I.F., Sander, C., Kirkwood, J.M., Kuchroo, V., and Zarour, H.M. (2010). Upregulation of Tim-3 and PD-1 expression is associated with tumor antigen-specific CD8^+^ T cell dysfunction in melanoma patients. Journal of Experimental Medicine 207, 2175–2186. 10.1084/jem.20100637.

39. Sakuishi, K., Apetoh, L., Sullivan, J.M., Blazar, B.R., Kuchroo, V.K., and Anderson, A.C. (2010). Targeting Tim-3 and PD-1 pathways to reverse T cell exhaustion and restore anti-tumor immunity. Journal of Experimental Medicine 207, 2187–2194. 10.1084/jem.20100643.

40. Zhou, Q., Munger, M.E., Veenstra, R.G., Weigel, B.J., Hirashima, M., Munn, D.H., Murphy, W.J., Azuma, M., Anderson, A.C., Kuchroo, V.K., and Blazar, B.R. (2011). Coexpression of Tim-3 and PD-1 identifies a CD8^+^ T-cell exhaustion phenotype in mice with disseminated acute myelogenous leukemia. Blood 117, 4501–4510. 10.1182/blood-2010-10-310425.

41. Grebinoski, S., Zhang, Q., Cillo, A.R., Manne, S., Xiao, H., Brunazzi, E.A., Tabib, T., Cardello, C., Lian, C.G., Murphy, G.F., et al. (2022). Autoreactive CD8^+^ T cells are restrained by an exhaustion-like program that is maintained by LAG3. Nature Immunology 23, 868–877. 10.1038/s41590-022-01210-5.

42. Workman, C.J., Cauley, L.S., Kim, I.-J., Blackman, M.A., Woodland, D.L., and Vignali, D.A.A. (2004). Lymphocyte Activation Gene-3 (CD223) Regulates the Size of the Expanding T Cell Population Following Antigen Activation In Vivo^1^. The Journal of Immunology 172, 5450–5455. 10.4049/jimmunol.172.9.5450.

43. Baitsch, L., Baumgaertner, P., Devêvre, E., Raghav, S.K., Legat, A., Barba, L., Wieckowski, S., Bouzourene, H., Deplancke, B., Romero, P., et al. (2011). Exhaustion of tumor-specific CD8^+^ T cells in metastases from melanoma patients. Journal of Clinical Investigation 121, 2350–2360. 10.1172/JCI46102.

44. Aine, M., Eriksson, P., Liedberg, F., Sjödahl, G., and Höglund, M. (2015). Biological determinants of bladder cancer gene expression subtypes. Scientific Reports 5, 10957. 10.1038/srep10957.

45. Wu, J., and Abraham, S.N. (2021). The Roles of T cells in Bladder Pathologies. Trends in Immunology 42, 248–260. 10.1016/j.it.2021.01.003.

