## Supplementary figures and tables for "SH2D2A is an indicator of favourable prognosis in bladder cancer and is enriched in activated Treg cells"

### Supplementary information

#### TABLE S1. Datasets used in this study.

| Name | Source | PMID | Type | Content |
|---|---|---|---|---|
| TCGA cancer dataset | TCGA (via HPA) | N/A | Bulk RNA-seq | Bulk RNA-seq of tumours from 21 cancer types |
| IMvigor210 | **`IMvigor210CoreBiologies`** (R package) | 29443960 | Bulk RNA-seq | Bulk RNA-seq of BLCA tumours and tumour metastases from patients undergoing αPD-L1 therapy |
| **`pbmcref`** | **`Azimuth`** (R package) | 34062119 | scRNA-seq + ADT assay | Reference PBMC dataset containing a 228-antibody ADT assay |
| GSE190888 | GEO | 35265520 | scRNA-seq | RNA-seq of bladder tissue from primary BLCA, recurrent BLCA, and cystitis glandularis |
| Tabula Sapiens bladder reference | CELL×GENE | 35549404 | scRNA-seq | RNA-seq of healthy human bladder tissue |
| GSE149652 | GEO | 32497499 | scRNA-seq | RNA-seq of T cells isolated from untreated BLCA, BLCA patients undergoing αPD-L1 therapy, and a BLCA patient undergoing chemotherapy |

#### TABLE S2. IMvigor210 metadata categories used in this study.

| Metadata name | Full name | Levels | Description |
|---|---|---|---|
| Lund type | Lund taxonomy classification[44] | Squamous cell carcinoma (SCC)-like | Type displaying particularly poor prognosis. Frequently TP53 mutated. Express markers for squamous differentiation, in addition to markers of basal cells of the urothelium. |
| | | Genomically unstable | Evenly split between high-grade non-muscle-invasive and muscle-invasive BLCA. Frequently TP53 mutated. Display genomic instability and high proliferation rates. |
| | | Infiltrated | Subgroup characterised by immune cell and stromal marker genes, indicative of a mixed-component tumour. |
| | | Urobasal A (UroA) | Predominantly low-grade papillary non-muscle-invasive tumours. Display frequent FGFR3 mutations and infrequent TP53 mutations. |
| | | Urobasal B (UroB) | Predominantly stage ≥ T1. Express markers of basal urothelium. TP53 mutated. |
| Immune phenotype | Tumour immune phenotype | Desert | No immune cells present |
| | | Excluded | Immune cells excluded from tumour proper |
| | | Inflamed | Active inflammation and immune penetration in tumour |
| | | Not recorded | Data not recorded |
| Enrollment IC score | Immune cell (IC) score at enrollment | IC0 | <1% immune cells intra- or peritumorally |
| | | IC1 | ≥1 to <5% immune cells intra- or peritumorally |
| | | IC2 | 5 to <10% immune cells intra or peritumorally |
| Binary αPD-L1 response | Response or lack of response to αPD-L1 therapy | CR/PR | Response to αPD-L1 therapy |
| | | SD/PD | No response to αPD-L1 therapy |
| Overall αPD-L1 response | Overall response to αPD-L1 therapy | CR | Complete response |
| | | PR | Partial response |
| | | SD | Stable disease |
| | | PD | Progressive disease |
| Survival | Survival | Yes | Patient survival to end of study |
| | | No | Patient not alive at end of study |

#### TABLE S3. CD4$^{+}$ cell type annotations from GSE149652 (Oh, *et al.*).

| Original annotation | Simplification | Description |
|---|---|---|
| $CD4_{CM}$ | CM | CD45RA$^{-}$ CCR7$^{+}$ central memory cells |
| $CD4_{CXCL13}$ | CXCL13$^{+}$ | CXCL13$^{+}$ IFNG$^{+}$ TOX$^{+}$ exhausted effector memory cells |
| $CD4_{TH17}$ | TH17 | IL17A$^{+}$ T helper 17 cells |
| $CD4_{ACTIVATED}$ | Activ. | CD69$^{+}$ FOXP3$^{-}$ activated effector cells |
| $CD4_{IL2RALO}$ | IL2RA$^{lo}$ | IL2RA$^{lo}$ FOXP3$^{+}$ TIGIT$^{+}$ TNFRSF4/9/18$^{+}$ CD27$^{+}$ Treg cells |
| $CD4_{IL2RAHI}$ | IL2RA$^{hi}$ | IL2RA$^{hi}$ FOXP3$^{++}$ TIGIT$^{+}$ TNFRSF4/9/18$^{++}$ CD27$^{+}$ Treg cells |
| $CD4_{GZMB}$ | GZMB$^{+}$ | GZMA$^{+}$ GZMB$^{++}$ NKG7$^{++}$ PRF1$^{+}$ GNLY$^{+}$ IFNG$^{+}$ cytotoxic cells |
| $CD4_{GZMK}$ | GZMK$^{+}$ | GZMA$^{+}$ GZMB$^{+}$ GZMK$^{++}$ NKG7$^{+}$ IFNG$^{+}$ cytotoxic cells |
| $CD4_{PROLIF}$ | Prolif. | MKI67$^{+}$ STMN1/TUBB$^{+}$ PCNA$^{+}$ HMGB1/2$^{+}$ proliferating cells |
| $CD4_{HSP}$ | HSP | Cells expressing various heat shock proteins |
| $CD4_{MITO}$ | Mito. | Cells expressing various mitochondria-linked genes |

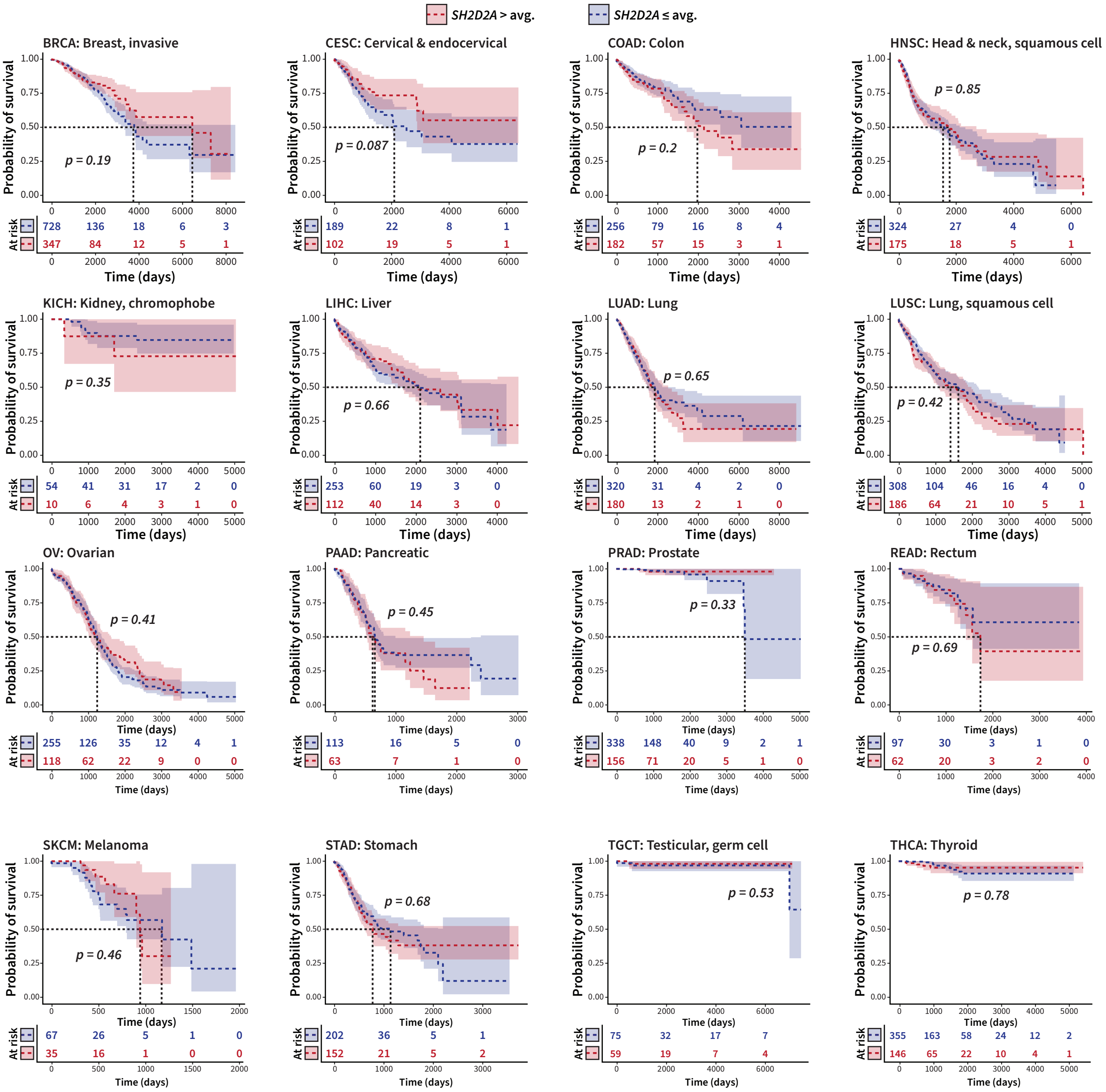


**Figure S1. Survival curves with no significant difference between SH2D2A high and low groups.** Kaplan-Meier survival curves of TCGA data of the 16 types of cancer with no statistically significant difference in survival between the SH2D2A > average (red) and the SH2D2A ≤ average (blue) populations.

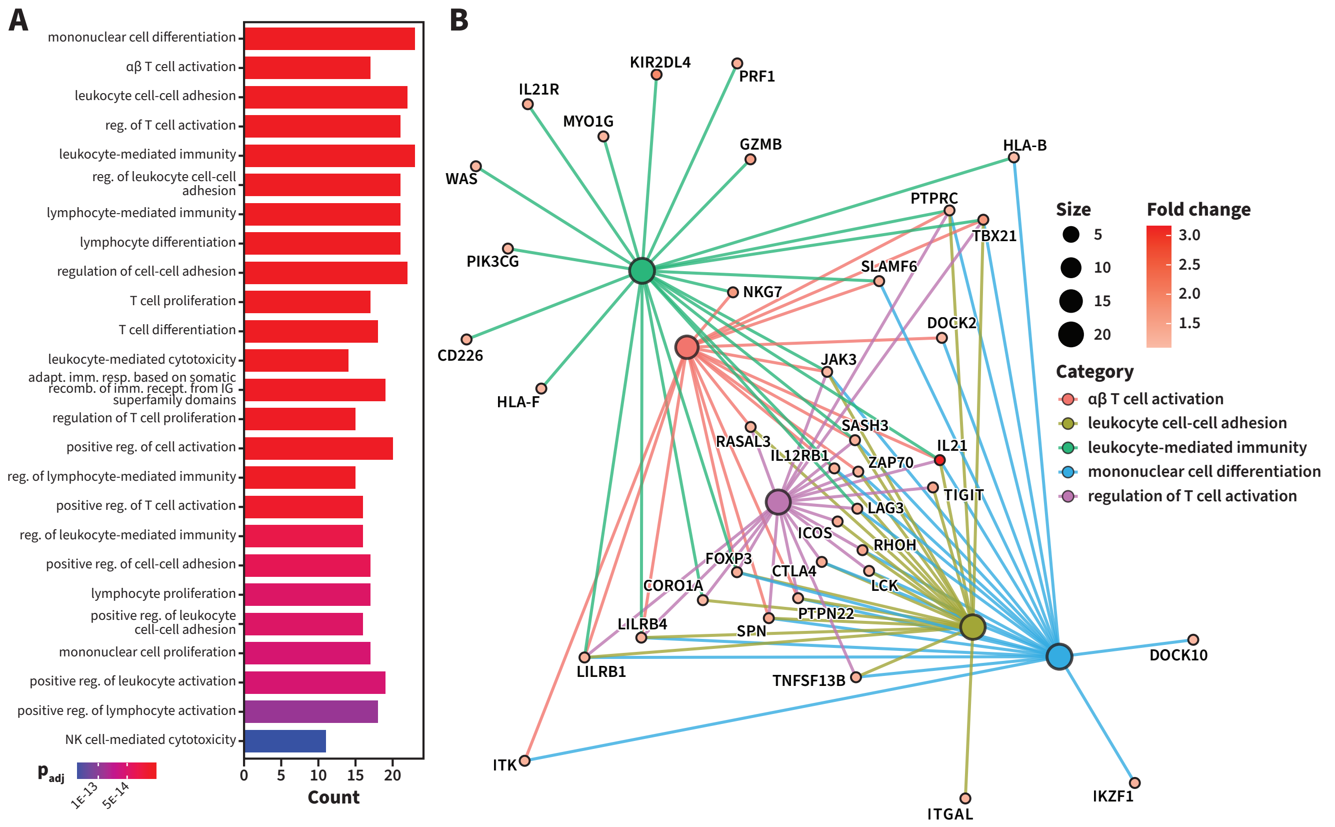


**Figure S2. GO pathway analysis of the 75 top-most genes shared in the IMvigor210 high SH2D2A group.** (A) Gene ontology (GO) pathway analysis of the top 25 upregulated biological processes within the set of the top 75 genes with the highest average Z-score in the high SH2D2A group from the IMvigor210 dataset, showing the gene counts for each GO result along with the adjusted p value of the enrichment. (B) Gene-concept network plot of the 5 most enriched GO results in the high SH2D2A group, showing the fold-changes of the involved genes as compared with the SH2D2A ≤ average cohort from the IMvigor210 dataset, as well as each gene's relation to the 5 GO categories.

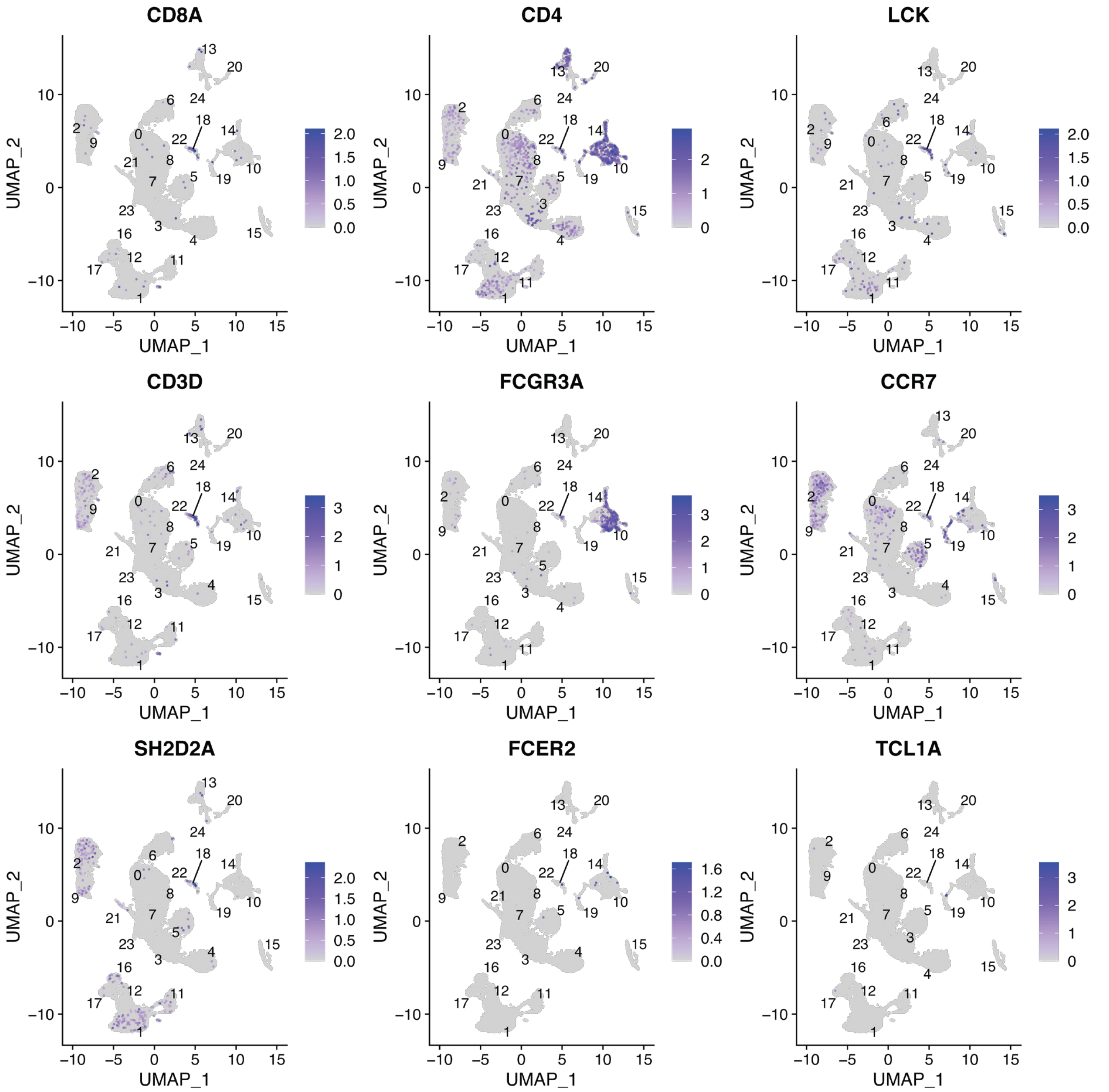


**Figure S3. Filtering of SH2D2A⁺ immune cells from GSE190888 with established immune cell markers.** Genes used to confirm the enrichment of SH2D2A in immune cells in the original full-tumour dataset of GSE190888. SH2D2A expression was largely confined to the small clusters 18 and 22. To offset the small cell number, clusters 1, 2, 9 and 11 were included in the initial selection based on limited expression of SH2D2A, CD4, and several other markers. The cells from these latter clusters were likely neoplastic and thus excluded from further analysis. Clusters 10 and 14 were candidates for containing immune cells due to high expression of CD4 and FCGR3A, but lacked expression of the critical kinase LCK, and were thus excluded.

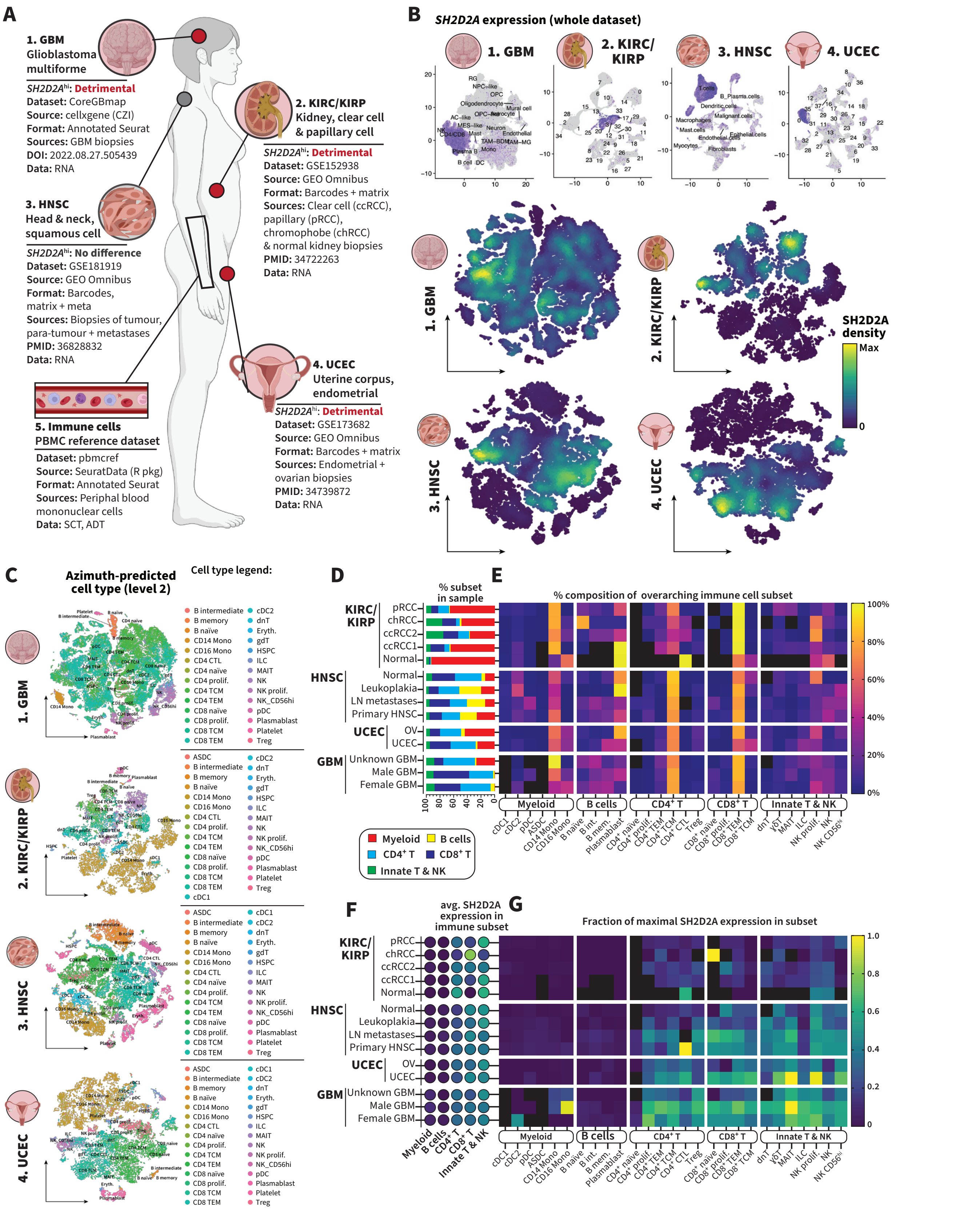

**Figure S4. Summary analysis of Azimuth-annotated cell types in scRNA datasets of GBM, UCEC, KIRP and KIRC.** (A) Graphical summary of the different scRNA datasets sourced for GBM (1), KIRC and KIRP (2), HNSC (3), and UCEC (4), along with their reference publications—additionally, the dataset pbmcref, used to annotate immune cell types via Azimuth, is included (5). (B) Initial UMAP plot for datasets 1-4 in (A) to isolate SH2D2A-enriched clusters and t-SNE plots of isolated SH2D2A-enriched clusters the whole datasets (as done for BLCA in Figure 4A). (D) Azimuth-derived cell type annotation of the SH2D2A-enriched clusters from the original datasets (as in Figure 4B). (D) Summary abundancies of the major immune cell subsets in each condition in each cancer subtype in (A) (as displayed for BLCA in Figure 4D, upper left). (E) Proportion of each Azimuth cell type annotation expressed as its fraction of the larger overarching immune cell subsets in (D) (as in Figure 4D, upper right). (F) Summary enrichment of SH2D2A in the major immune cell subsets in each condition in each cancer subtype in (A) (as in Figure 4D, lower left). (G) Proportion of SH2D2A expression represented by each Azimuth cell type annotation expressed as a fraction of the large immune cell subset's total SH2D2A expression (as in Figure 4D, lower right).
